# Glucose promotes neuron morphology abnormalities and egg-laying defects dependent of the serotonin signaling pathway in *Caenorhabditis elegans*

**DOI:** 10.1101/2025.09.20.677429

**Authors:** Manuel A. Ruiz, Dennis R. Dumesnil, Abhishek Shah, Mary L. Ladage, Luhua Song, Pamela A. Padilla

**Author notes:** Correspondence to: Pamela A. Padilla.

## Abstract

A chronic hyperglycemic state can lead to neuropathological complications such as permanent nerve damage. Neuropathology is a debilitating ailment that affects nearly half of the individuals with diabetes, underscoring the relevance to investigate the impact of diet and glucose levels on the nervous system. Using the genetic model system *Caenorhabditis elegans*, we determined that a diet rich in glucose impacts biological processes dependent on neuronal function. The rate of intrauterine egg-hatching increased in animals fed a glucose diet and the inhibition of dopamine or serotonin is sufficient to counteract the impact of the glucose diet. The glucose-induced intrauterine egg-hatching phenotype is independent of the SER-1 and SER-7 receptors yet dependent on the DAF16/FOXO transcription factor. Furthermore, animals fed a glucose-supplemented diet displayed abnormal morphologies in the HSN and NSM serotonergic neurons and the HSN displayed axonal degeneration, which is a prominent pathological feature in neurological dysfunction and disease. Combined these data indicate that a glucose-diet affects nervous system and signaling pathways in *C. elegans*, which can be used to further model diet-induced neuropathy.

## INTRODUCTION

A high glycemic index diet increases the risk of obesity, stroke, cardiovascular disease, and diabetes. The number of people with type 2 diabetes has substantially increased in the last four decades, from 108 million people in the 1980s to 422 million people in 2014 and within the U.S., 37.3 million people (11.3%) have diabetes (Diabetes Research Institute, 2023) (Roglic and World Health Organization 2016; Herman *et al*. 2023). Diabetes is a contributing factor or cause for many health conditions including stroke, kidney failure, blindness, heart attack, and lower limb amputation. Diabetes-induced peripheral neuropathy is observed in about 50% of diabetic adults and is associated with chronic pain, lower limb ulcers, lower limb amputation, and an increase in morbidity (Pop-Busui *et al*. 2022). Although the negative impact of diabetic neuropathy is understood, the mitigation of such is challenging.

The nematode *Caenorhabditis elegans* is a well-established genetic model system used to understand neural circuitry and behavior (Sengupta and Samuel 2009). The *C. elegans* nervous system is made up of 302 neurons, which is nearly a third of the total cells found in the nematode (wormatlas.org, 2023). Furthermore, like humans, *C. elegans* have neurotransmitters that modulate sensory and motor information. Among the most studied neurotransmitters in *C. elegans* are the biogenic amines dopamine and serotonin, which modulate behaviors such as learning, locomotion, pharyngeal pumping, mating behaviors, and egg-laying (Loer and Kenyon 1993; Schafer and Kenyon 1995; Waggoner *et al*. 1998; Duerr *et al*. 1999; Nurrish *et al*. 1999; Sawin *et al*. 2000; Sze *et al*. 2000). In the case of egg-laying, some studies indicate that dopamine inhibits egg-laying (Schafer and Kenyon 1995; Weinshenker *et al*. 1995; Sanyal *et al*. 2004), whereas the release of serotonin, by the hermaphrodite specific motor neurons (HSN) that innervate the vulva muscles, stimulates egg laying (Waggoner *et al*. 1998). The role of specific neurotransmitters in response to diet could lead to deeper understanding metabolic diseases in humans. A glucose-supplemented diet is thought of as a dietary stress since it is associated with reduced brood size, reduced lifespan, sensitivity to oxygen deprivation, and increase in stress response transcripts (Lee *et al*. 2009; Garcia *et al*. 2015). In addition, glucose reduces egg-laying rate and increases egg retention and internal hatching (Teshiba *et al*. 2016; Alcantar-Fernandez *et al*. 2018).

In humans, serotonin regulates a diverse range of behaviors and functions such as sleep, mood, wound healing, blood clotting, sexual desire, learning, and cognition. Serotonergic dysfunction is linked to the incidence of obsessive-compulsive disorder, anxiety, depression, Down syndrome, autism spectrum disorder, schizophrenia, and obesity (Whitaker-Azmitia 2001). In the human body, peripheral serotonin is synthesized in the enterochromaffin cells and circulating serotonin is mostly found stored in platelets (Cai *et al*. 2022). In the central nervous system, serotonin is mostly synthesized in neurons originating in the raphe nuclei and functions as a neurotransmitter. Recent studies have associated a link between serotonin and diabetes. Tryptophan hydroxylase inhibitors are potential targets to treat diabetes and obesity (Kolodziejczak *et al*. 2015; Abg Abd Wahab *et al*. 2019). Yet further studies are needed to gain a greater understanding of the relationship between metabolic dysfunction, diabetes, and serotonin signaling.

In this study, we demonstrate that in *C. elegans* a glucose-supplemented diet increases the proportion of intrauterine egg-hatching (IUEH), a phenomenon in which embryos are abnormally retained and hatch inside the uterus. A glucose-supplemented diet upregulates the expression of the *cat-2* and *tph-1* genes responsible for the synthesis of the rate-limiting enzymes for dopamine and serotonin production. The dopamine and serotonin antagonist drugs chlorpromazine and methiothepin respectively, suppress the glucose induced IUEH phenotype in wildtype hermaphrodites. In addition, a glucose diet impacts the morphology of two serotonergic neurons, the HSN and the neurosecretory motor (NSM) neurons, and increases the incidence of HSN axonal degeneration in adult animals. Together these results provide a greater insight into the link between glucose, serotonin, and organ dysfunction which could elucidate our understanding of disease states such as diabetes.

## RESULTS

### Impact of a glucose-supplemented diet on egg-laying

The production of offspring can be impacted by the environment and in *C. elegans,* mating or starvation can induce intrauterine egg hatching and matricide phenotypes in wild-type (N2) hermaphrodite animals (Chen and Caswell-Chen 2003; Pickett and Kornfeld 2013). While examining N2 hermaphrodites raised on a glucose-supplemented diet, we observed a significant increase in the number of animals with the intra<u>u</u>terine <u>e</u>gg <u>h</u>atching (IUEH) and matricide phenotype, relative to animals raised on a standard OP50 diet (Figure 1A). The IUEH phenotype induced by a glucose diet was observed in day 3 to day 10 adult hermaphrodites (Figure 1B). Thus, in further experiments, animals were assayed until day 10 of adulthood. To determine if the glucose diet impacted developing and/or adult animals, we examined animals fed a glucose-supplemented diet only during larval development or only during adulthood. Animals assayed include those raised on the: control OP50 diet from L1 larvae through adulthood (G−); glucose-supplemented diet from L1 larvae through adulthood (G+); glucose-supplemented diet during larval development and then fed a standard diet through adulthood (G+ to G−); standard OP50 diet during larval development and then fed a glucose-supplemented diet through adulthood (G− to G+) (Figure S1). We determined that animals fed a glucose-supplemented diet during larval development had a significant increase in the IUEH phenotype relative to animals only fed a control diet throughout life (G+ to G−, Figure 1C, D). In contrast animals fed a control diet during larval development but a glucose-supplemented diet through adulthood did not have a significant increase in the IUEH phenotype relative to animals only fed a control diet throughout life (G− to G+, Figure 1C, D). Furthermore, the animals fed a glucose-supplemented diet during larval development (G+ to G−) exhibited a reduced brood size (Figure 1E) and egg-laying rate relative to control (Figure 1F). Whereas the animals fed a glucose-supplemented diet only through adulthood (G− to G+) did not have a significant difference in brood size or number of eggs laid relative to control (Figure 1E, F, respectively). These data indicate that glucose-supplementation during larval development and not throughout adulthood impacts egg-laying and offspring production. This aligns with previous findings that a glucose-supplemented diet impacts brood size and egg-laying rate (Lee *et al*. 2009; Tauffenberger and Parker 2014; Teshiba *et al*. 2016; Alcantar-Fernandez *et al*. 2018).

**Figure 1.**
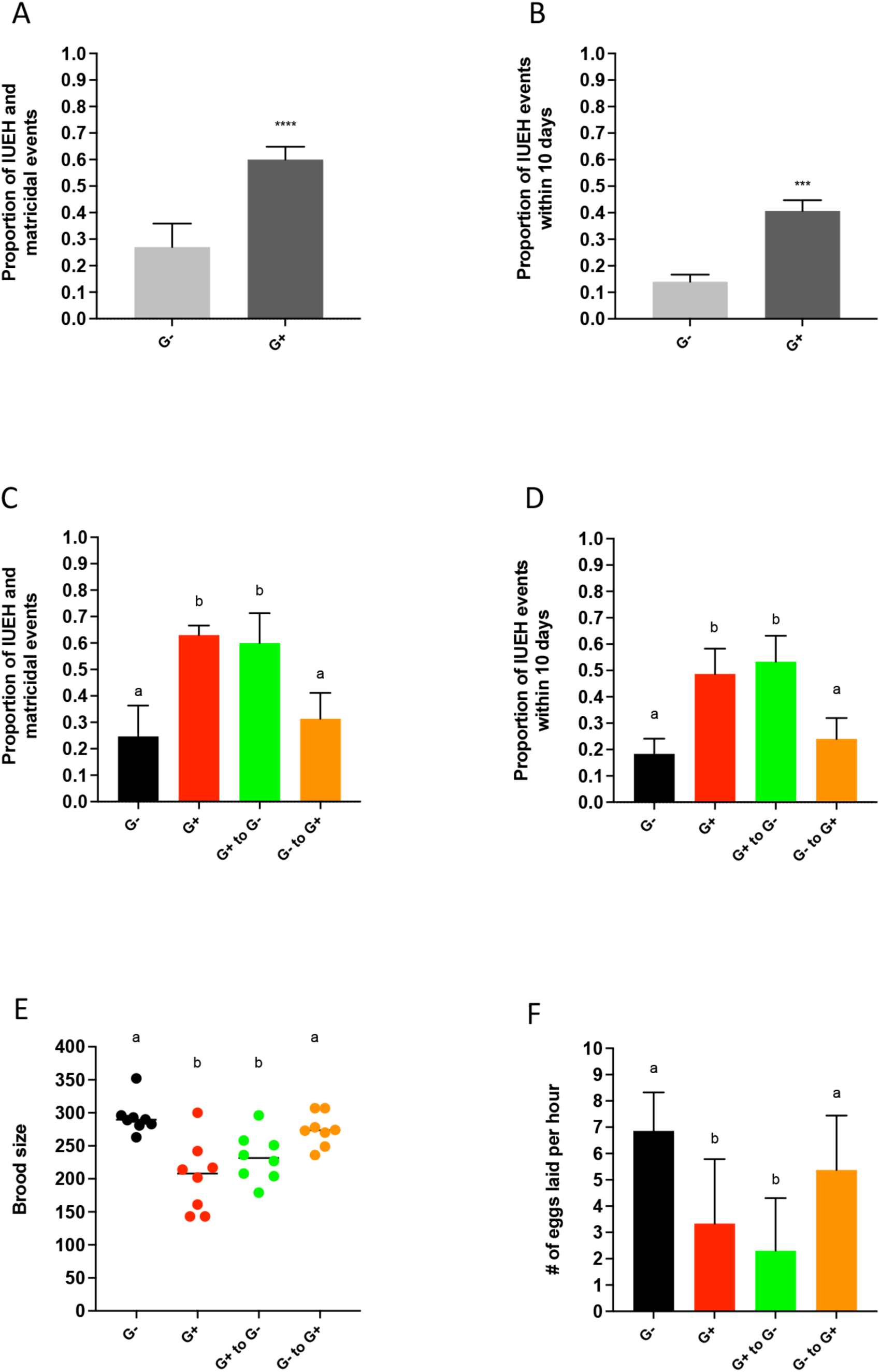
A glucose-supplemented diet impacts the production of offspring. Feeding regimen includes: N2 hermaphrodites fed a glucose-supplemented diet from L1 to indicated adult stage (G+), control diet of OP50 *E. coli* from L1 larval stage through indicated adult stage (G−), animals fed a glucose-supplemented diet during larval development and a control diet throughout adulthood (G+ to G−), animals fed a control diet during larval development and a glucose-supplemented diet through adulthood (G− to G+). (A, B) The proportion of intrauterine egg hatching and matricide (IUEH) was assayed daily in animals (A, C) throughout their lifetime, or (B, D) during the first ten days of adulthood. (A, B) two-tailed unpaired t-test with at least three independent experiments were conducted (n>150); (A) **** indicates p < 0.0001; (B) *** indicates p < 0.01. (E) The total brood size or (F) the number of eggs laid per hour by 1-day old hermaphrodites was determined. For C-F, identical letters represent groups with no significant difference and non-identical letters represent groups with significant difference (one-way ANOVA, Tukey’s multiple comparisons, p < 0.05). At least three independent experiments with n>150 (C, D), n>30 (E), n=8 (F).

### Reduction in dopamine synthesis suppresses the glucose induced IUEH phenotype

Previously we identified 2,370 genes that are differentially expressed in wild-type non-gravid adult animals fed a glucose-supplemented diet (Garcia *et al*. 2015). Included in this dataset are transcripts that code for proteins that are thought to have nervous system function (17 downregulated transcripts and 133 upregulated transcripts, S1 Table). One of the genes upregulated in glucose-fed animals is *cat-2*, which encodes a putative tyrosine hydroxylase essential for the synthesis of dopamine and inhibiting egg laying in *C. elegans* (Schafer and Kenyon 1995). We confirmed that there is a significant increase in *cat-2* expression, in non-gravid adult hermaphrodites fed a glucose-supplemented diet relative to those fed standard OP50 diet, as detected using qRT-PCR; this verifies the RNA-sequencing data (Figure 2A). Thus, we investigated the glucose-induced IUEH phenotype in animals with a reduced level of dopamine due to genetic mutation that affects dopamine synthesis. The *cat-2(n4547)* and *bas-1(tm351)* mutants were examined; *bas-1* is predicted to encode the dopamine synthesis protein L-dopa decarboxylase. Unlike N2 animals fed a glucose-supplemented diet, both the *cat-2(n4547)* and *bas-1(tm351)* animals fed a glucose-supplemented diet did not have a significant increase in the IUEH phenotype relative to respective animals fed an OP50 control diet (Figure 2B, C). To further assess the role of dopamine in the glucose-induced IUEH phenotype, we examined N2 glucose-fed animals treated with a dopamine antagonist (chlorpromazine, 1.5 µM) or exogenous dopamine (56 µM)￼(Loer and Kenyon 1993; Waggoner *et al*.￼. Chlorpromazine treatment suppressed the glucose-induced IUEH phenotype in wild-type animals (Figure 2D). However, exogenous dopamine supplementation was not sufficient to increase the IUEH phenotype relative to respective controls (Figure 2E, F). Together, these results indicate that a reduction of dopamine in N2 animals suppresses the glucose-induced IUEH phenotype, but exogenous feeding of dopamine was not sufficient to induce the IUEH phenotype in N2 animals.

**Figure 2.**
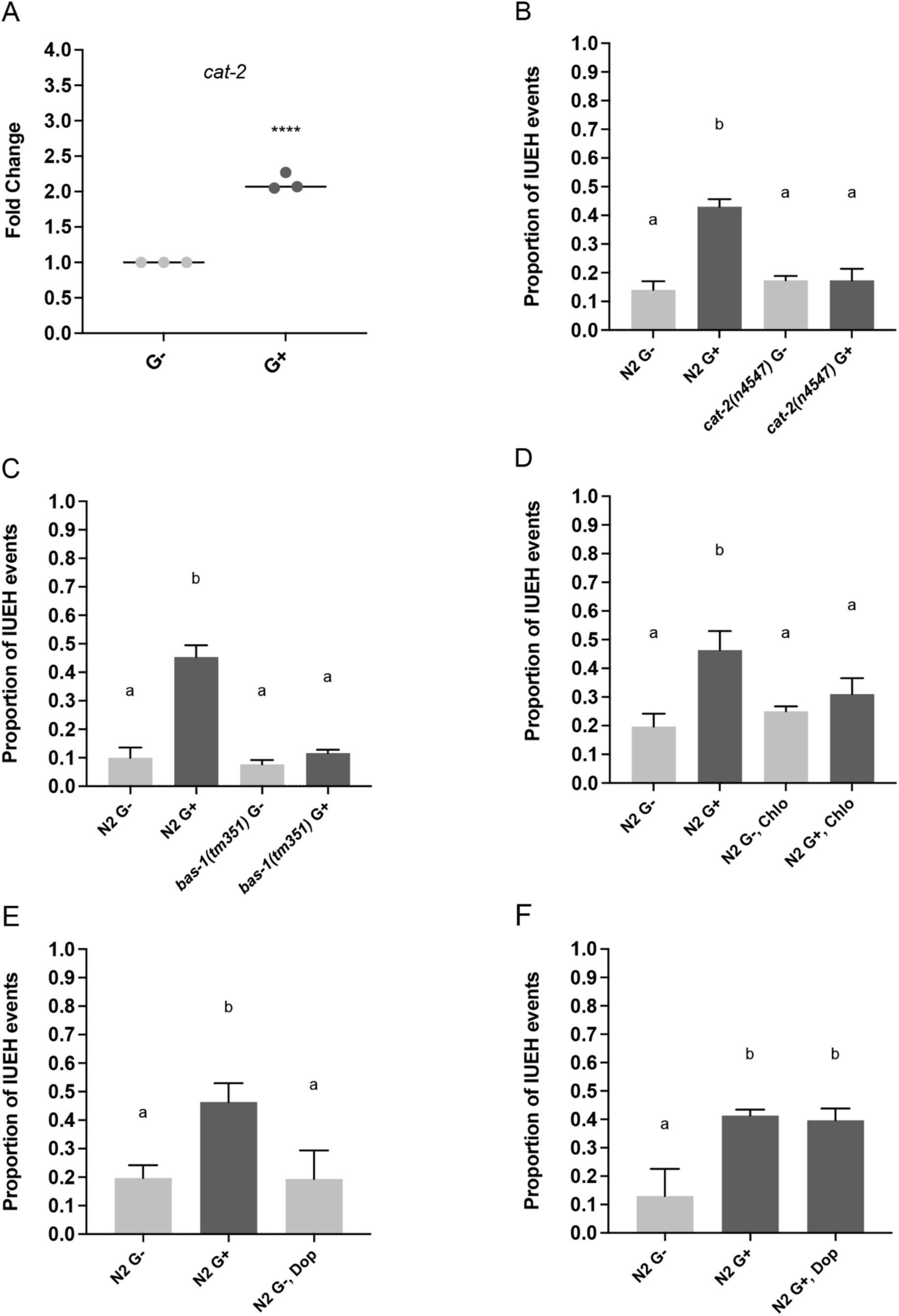
The reduction of dopamine synthesis suppresses the glucose-induced intrauterine egg-hatching (IUEH) phenotype. (A) qRT-PCR experiment to determine the fold change of *cat-2* expression in wild-type animals fed a control or a glucose-supplemented diet. The *cat-2* expression is significantly increased in the N2 non-gravid young adult fed a glucose-supplemented diet. The **** indicates p < 0.0001; two-tailed unpaired t-test; at least three independent experiments were conducted with >80 animals per experiment for a total of n>240 animals. The proportion of animals displaying the IUEH phenotype was determined in animals fed an OP50 glucose-supplemented diet (G+) or control OP50 diet (G−) in (B) *cat-2(n4547)* animals, (C) *bas-1(tm351)* animals, (D) N2 animals provided a dopamine antagonist (1.5 µM chlorpromazine, Chlo), and (E, F) N2 animals provided with exogenous dopamine (56 µM). (B-F) Animals were examined for the IUEH phenotype during the 10 days of adulthood. Identical letters represent groups with no significant difference. Non-identical letters represent groups with significant difference (p < 0.01, one-way ANOVA, Tukey’s multiple comparisons; at least three independent experiments were conducted with n > 150).

### Serotonin is necessary and sufficient for the glucose-induced IUEH phenotype

Egg-laying behavior in *C. elegans* is modulated by hermaphrodite specific motor neurons (HSNs) and various studies indicate that the HSNs utilize serotonin (Horvitz *et al*. 1982; Weinshenker *et al*. 1995). Notably, exogenous serotonin rescues egg-laying defects in animals with ablated HSNs (Trent *et al*. 1983). To examine if serotonin affects the glucose-induced IUEH phenotype, we exposed glucose-fed animals to exogenous serotonin (56 µM 5-HT) or the serotonin antagonist methiothepin (1.5. µM) N2 animals fed a glucose-supplemented diet and exogenous serotonin had a significant increase in the IUEH phenotype relative to animals fed just the glucose-supplemented diet (Figure 3A). This indicates that the addition of exogenous serotonin intensifies the effect of a glucose diet. The animals treated with the serotonin antagonist (1.5 µM methiothepin) showed a significant decrease in the IUEH phenotype relative to animals fed just the glucose-supplemented diet (Figure 3B). We observe that N2 animals fed an OP50 diet and exogenous serotonin (56 µM 5-HT) have a significant increase in the IUEH phenotype relative to the N2 animals fed OP50 (Figure 3B), which aligns with previous reports (Schafer and Kenyon 1995).

**Figure 3.**
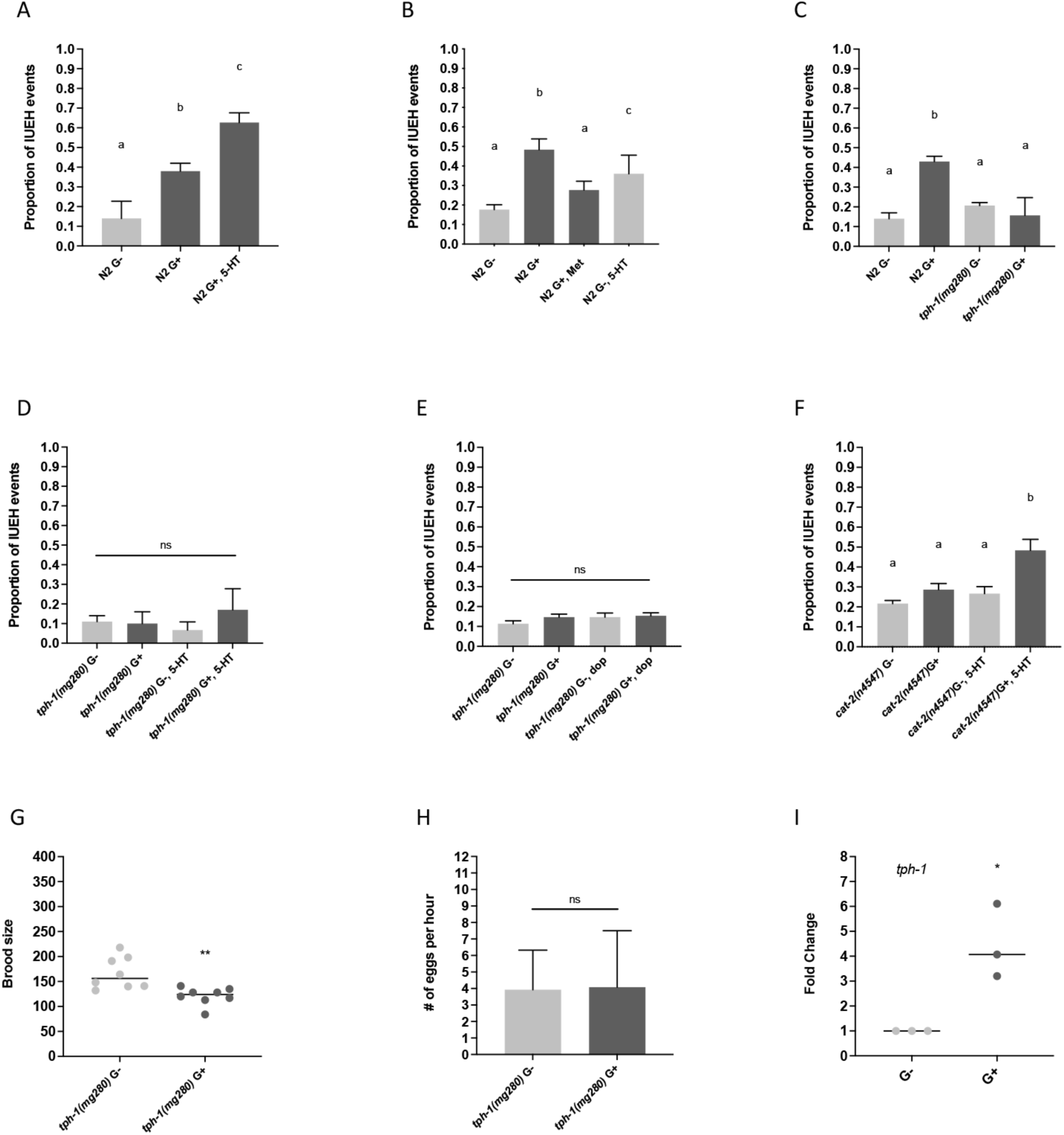
The glucose-induced IUEH phenotype is dependent on serotonin. (A-F) The IUEH phenotype was examined in the specified strains and diets. (A) N2 animals fed either a glucose-supplemented diet (G+) with or without exogenous serotonin (56 µM 5-HT). (B) N2 animals fed a standard OP50 diet (G−) with or without exogenous serotonin (56 µM 5-HT); a glucose-supplemented diet (G+) with or without the serotonin inhibitor methiothepin (1.5 µM Met). (C) The *tph-1(mg280)* animals fed a glucose-supplemented diet (G+) or standard OP50 diet (G−). (D) The *tph-1(mg280)* animals fed a glucose-supplemented diet (G+) or standard OP50 diet (G−) with or without exogenous serotonin (56 µM 5-HT). (E) The *tph-1(mg280)* animals fed a glucose-supplemented diet (G+) or standard OP50 diet (G−) with or without exogenous dopamine (G+, 56 µM Dop). (F) The *cat-2(n4547)* animals fed a glucose-supplemented diet (G+) or standard OP50 diet (G−) with or without exogenous serotonin (56 µM 5-HT). (A-F) Identical letters represent groups with no significant difference. Non-identical letters represent groups with significant difference p < 0.01 and no significant difference is indicated by ns; (one-way ANOVA, Tukey’s multiple comparisons; at least three independent experiments were conducted with n > 150). Animals were examined for the phenotype during the first 10 days of adulthood. (G) Brood size was assayed in the *tph-1(mg280)* animals fed either a glucose diet (G+) or a standard diet (G−). The * indicates p < 0.05, two-tailed unpaired t-test; three independent experiments with a total of n=8. (H) The number of eggs laid per hour was determined for the *tph1(mg280)* 1-day old adult animals fed either a glucose diet (G+) or a standard OP50 diet (G−). (I) qRT-PCR experiment to determine the fold change of *tph-1* in wild-type N2 young non-gravid adult animals fed either a glucose-supplemented diet or standard OP50 diet were assayed. The * indicates p < 0.05; two-tailed unpaired t-test; at least three independent experiments were conducted with >80 animals per experiment for a total of n>240 animals.

Tryptophan hydroxylase (TPH-1) is required for the biosynthesis of serotonin (Sze *et al*. 2000). We examined the *tph-1(mg280)* mutant and determined that TPH-1 function is required for the glucose-induced IUEH phenotype (Figure 3C). Furthermore, the feeding of exogenous serotonin (56 µM 5-HT) to the *tph-1(mg280)* mutant, fed a standard OP50 diet or a glucose-supplemented diet, was not sufficient to significantly increase the IUEH phenotype (Figure 3D). It is also the case that the feeding of exogenous dopamine is not sufficient to induce the glucose-induced IUEH phenotype in *tph-1(mg280)* animals (Figure 3E). Yet, the *cat-2(n4547)* animal fed exogenous serotonin and a glucose-supplemented diet had a significant increase in the IUEH phenotype (Figure 3F). Combined, these data indicate serotonin is necessary and sufficient for the glucose-induced IUEH phenotype in N2 animals, and that functional TPH-1, but not CAT-2, is necessary to observe the compounding effect of exogenous serotonin in animals fed a glucose-supplemented diet.

We next examined how *tph-1* dysfunction impacts offspring production in animals fed a glucose-supplemented diet. Like N2 animals (Figure 1G), the *tph-1(mg280)* animals fed a glucose-supplemented diet had a decrease in the brood size relative to those fed a standard diet (Figure 3G). However, a glucose diet did not significantly decrease the egg laying rate in the *tph-1(mg280)* animals (Figure 3H). Through qRT-PCR analysis we found that the *tph-1* transcript increases in animals fed a glucose diet (Figure 3I) further supporting the idea that serotonergic processes respond to a glucose diet. An additional indicator of dysfunctional egg-laying is the presence of older embryos within the uterus of the adult animal. We observed that, in comparison to control, N2 animals fed a glucose-supplemented diet hold embryos within the uterus that are at or past the 1.5-fold embryonic stage (5-day old adult animal, Figure S2A, B). However, this phenotype was not observed in *tph-1(mg280)* adult animals (Figure S2C, D). Combined, this aligns with previous findings that serotonergic signaling is required to observe the glucose-induced egg-laying defect (Teshiba *et al*. 2016).

### The glucose-induced IUEH phenotype is independent of the SER-1 or SER-7 receptor

If serotonin is required for the glucose-induced IUEH phenotype, then it is plausible that disruption of the predicted serotonin receptor(s) would have an impact on the phenotype. SER-1 and SER-7, part of the serotonin/octopamine receptor family, are expressed in various tissues including the vulva muscles and have a role in egg-laying in N2 hermaphrodites (Carnell *et al*. 2005; Dempsey *et al*. 2005; Hobson *et al*. 2006) (Figure 4A). The *ser-1(ok345)* animals fed a glucose diet exhibited the IUEH phenotype, and exposure to serotonin (56 µM 5-HT) did not significantly alter the IUEH phenotype, relative to the representative control (Figure 4B). This indicates that the feeding of a glucose-supplemented diet is the primary determinant of the IEUH phenotype in the *ser-1(ok345)* animal. The *ser-7(tm1325)* animal is known to have an egg-laying defect (Hobson *et al*. 2006; Xiao *et al*. 2006). We observe that the *ser-7(tm1325)* animal and the *ser-1(ok345);ser-7(tm1325)* double mutant animal exhibited the IUEH phenotype when fed either the standard OP50 diet or a glucose-supplemented diet (Figure 4C, D). The presence of glucose and serotonin (56 µM 5-HT) leads to a significant increase in the IUEH phenotype relative to the animals only exposed to serotonin (56 µM 5-HT) or fed glucose (Figure 4C, D). Recall that the serotonin antagonist (1.5 µM methiothepin) suppressed the glucose induced IUEH phenotype in N2 animals (Figure 3B). However, the serotonin antagonist (1.5 µM methiothepin) was unable to suppress the IUEH phenotype observed in the *ser-1(ok3456)* animal fed a glucose diet (Figure 4E) or the *ser-7(tm1325)* animal fed either a standard or glucose diet (Figure 4F). Combined, this data suggests that the glucose-induced IUEH phenotype does not require SER-1 or SER-7 function and that serotonin may act on a receptor(s) besides SER-1 or SER-7 to induce the IUEH phenotype. Thus, unlike the disruption of serotonin synthesis, the alteration of serotonin signaling via SER-1 and/or SER-7 dysfunction does not suppress the glucose-induced IUEH phenotype. Therefore, it is possible that methiothepin suppresses the glucose-induced IUEH phenotype via SER-1 and SER-7 loss produces a “ceiling effect” that is independent of a glucose-supplemented diet and methiothepin.

**Figure 4.**
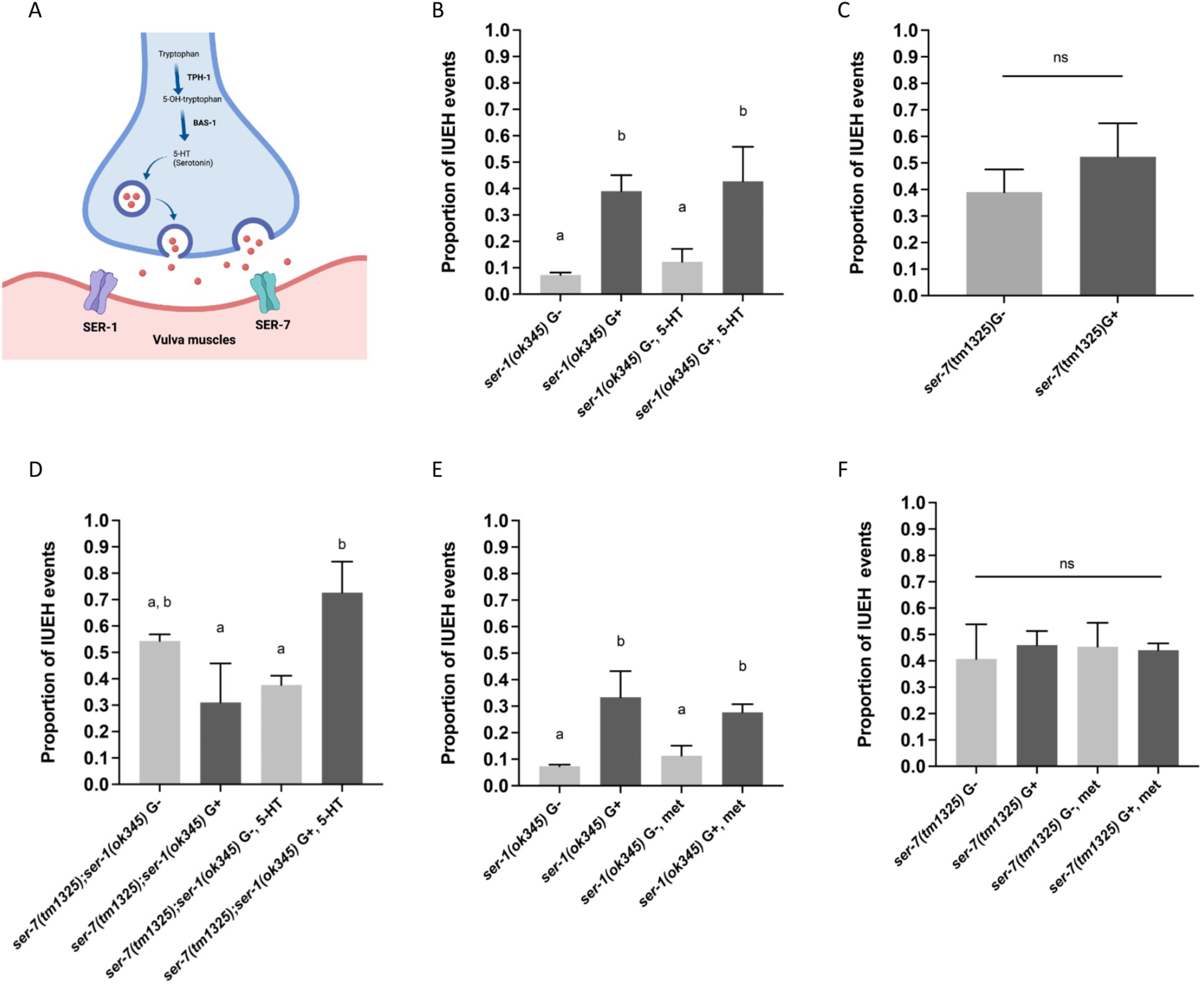
The glucose-induced intra-uterine egg hatching phenotype is independent of the SER-1 or SER-7 receptor. (A) Illustration of the hermaphrodite-specific neuron (HSN) serotonergic neuromuscular junction in *C. elegans*. The IUEH phenotype was examined in (B) *ser-1(ok345)*; (C) *ser-7(tm1325)*; or (D) *ser-7(tm1325);ser-1(ok345)* animals fed either a standard or glucose-supplemented diet, in the absence or presence of exogenous serotonin (56 µM 5-HT). The IUEH phenotype was examined in (E) *ser-1(ok345)* or (F) *ser-7(tm1325)* animals fed either a standard or glucose-supplemented diet in the absence or presence of methiothepin, the serotonin inhibitor (1.5 µM met). (B-F) Non-identical letters represent groups with significant difference p value < 0.05 (one-way ANOVA, Tukey’s multiple comparisons; at least three independent experiments were conducted with n > 150). Animals were examined during the first ten days of adulthood.

### The glucose-induced IUEH phenotype is dependent on DAF-16/FOXO

The DAF-16/FOXO transcription factor functions downstream of the DAF-2 receptor/DAF-18 phosphatase. Dysfunction of the insulin-like signaling DAF-16/FOXO transcription factor leads to defects in the HSN migration during development (Kennedy *et al*. 2013) and phenotypes associated with a glucose-supplemented diet are dependent on the DAF-16/FOXO transcription factor (Lee *et al*. 2009; Garcia *et al*. 2015). We tested if the glucose-induced IUEH phenotype was impacted in animals with altered insulin-like signaling. Interestingly, the glucose-induced IUEH phenotype was not observed in the *daf-16(mu86)* (Figure 5A). The addition of exogenous serotonin (56 µM 5-HT) induced the IUEH phenotype in the glucose-fed *daf-16(mu86)* animal but not those animals fed a control OP50 diet (Figure 5A). The IUEH phenotype is observed in the *daf-2(e1370)* animal fed a standard OP50 food or a glucose-supplemented diet and the feeding of exogenous serotonin (56 µM) or the serotonin antagonist (methiothepin, 1.5 µM) did not significantly alter the IUEH phenotype in the *daf-2(e1370)* animal fed a standard or glucose-supplemented diet (Figure 5B). However, the *daf-16(mu86)* mutation reduced the IUEH phenotype in the *daf-2(e1370)* animals fed a standard OP50 diet but not a glucose-supplemented diet (Figure 5C). This suggests that the IUEH phenotype can be modulated by both glucose and *daf-2* function. Unlike wildtype animals, a glucose-supplemented diet does not reduce the brood size or number of eggs laid per hour in the *daf-16(mu86)* animals (Figure 5D, E). There is a slight but significant decrease in brood size (Figure 5F) but not the number of eggs laid per hour in the *daf-2(e1370)* animal fed a glucose diet (Figure 5G). Combined, these data indicate that DAF-16 regulates the serotonin response to a glucose-supplemented diet, is required for the glucose-induced phenotypes, and promotes the IEUH phenotype in the *daf-2(e1370)* animal.

**Figure 5.**
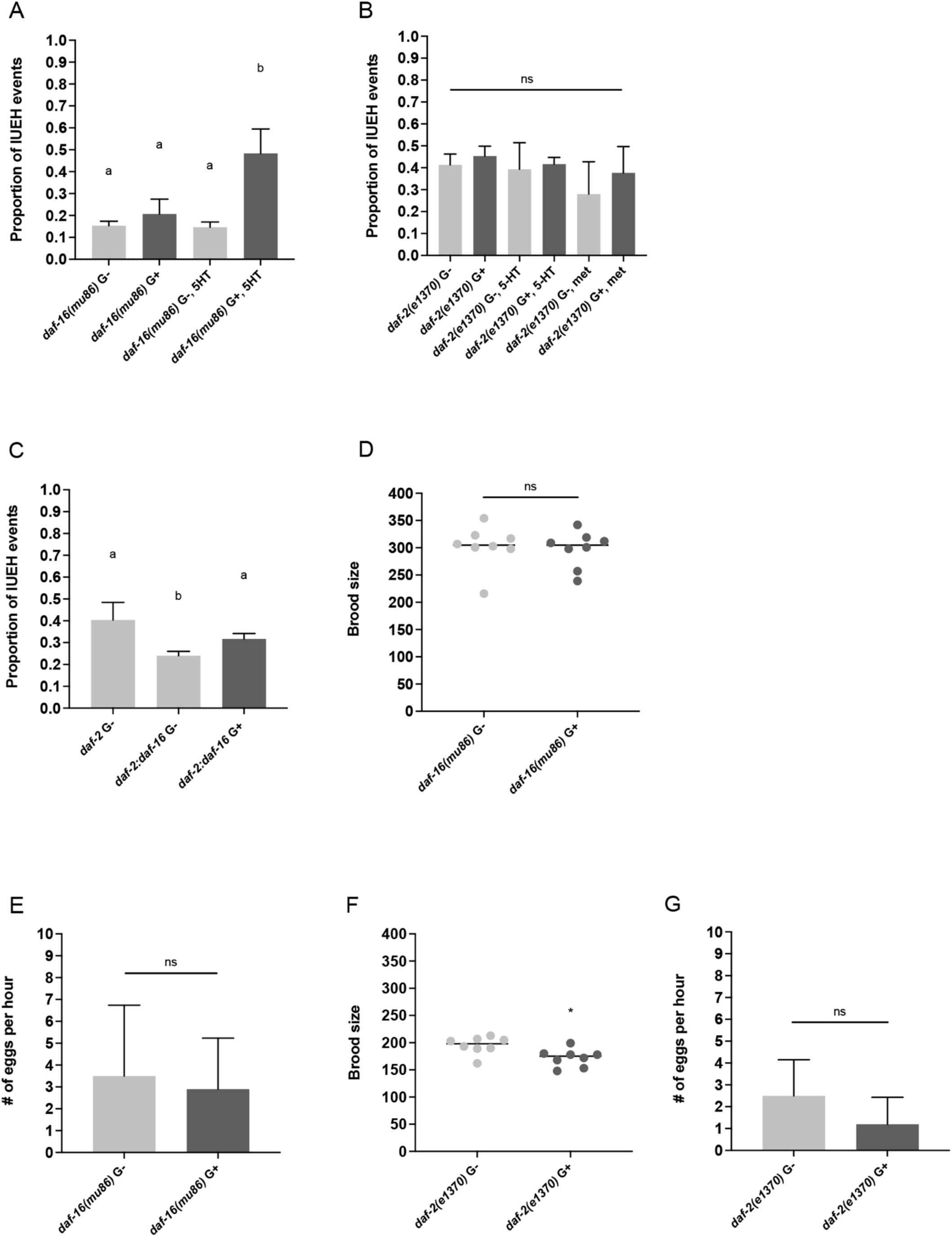
DAF-16 is required for glucose-induced phenotypes. (A-C) The IUEH phenotype was examined in the specified strains and diet during the first ten days of adulthood. (A) The *daf-16(mu86)* animal fed either a glucose-supplemented diet (G+) or standard OP50 diet (G−) with or without exogenous serotonin (56 µM 5-HT). (B) The *daf-2(e1370)* animal fed either a glucose-supplemented diet (G+) or standard OP50 diet (G−) in the presence or absence of exogenous serotonin (56 µM 5-HT) or methiothepin, the serotonin inhibitor (1.5 µM met). (C) The *daf-2(e1370);daf-16(mu86)* animal fed either a glucose-supplemented diet (G+) or standard OP50 diet (G−). (A-C) The non-identical letters represent groups with significant difference p value <0.001 (one-way ANOVA, Tukey’s multiple comparisons; at least three independent experiments were conducted with n > 150). The brood size and number of eggs laid per hour was determined in *daf-16(mu86)* animals (D, E, respectively) or *daf-2(e1370)* animals (F, G, respectively) fed either a glucose-supplemented diet (G+) or standard OP50 diet (G−). (D, E) The * indicates p < 0.05; ns indicates not significant (two-tailed unpaired t-test). At least three independent experiments were conducted with n>150.

### A glucose-supplemented diet impacts HSN neuronal morphology

The serotonergic HSNs are essential for normal egg-laying and have an oval-like shape present near the vulva region with axons extending towards the posterior region of the animal (Altun 2025). Using the *tph-1::GFP* transcriptional reporter to visualize the HSN in adult animals fed a standard OP50 diet (G−) or glucose-supplemented diet (G+), we found that the HSN is morphologically abnormal in animals fed a glucose-supplemented diet (Figure 6A-C). We verified that animals fed a glucose-supplemented diet had an increase in abnormal HSN morphology using another reporter, *ida-1::GFP*, which is expressed in the HSNs (Figure S3A). The glucose-induced HSN abnormalities include abnormal positioning of the HSN, extending towards near the dorsal region (Figure 6A, yellow arrow), and abnormal branching, axonal growths, and ectopic somas (Figure 6A, yellow arrows). Furthermore, in comparison to animals fed a standard OP50 diet, animals fed a glucose-supplemented diet also exhibited axonal degeneration, as determined by discontinuous GFP labeling (Figure 6D-F). The feeding of exogenous 5-HT did not increase the abnormal glucose-induced HSN morphology (Figure 6G, H). However, the serotonin inhibitor (methiothepin) reduced the abnormal HSN phenotype in 3-day old animals fed a glucose diet (Figure 6H).

**Figure 6.**
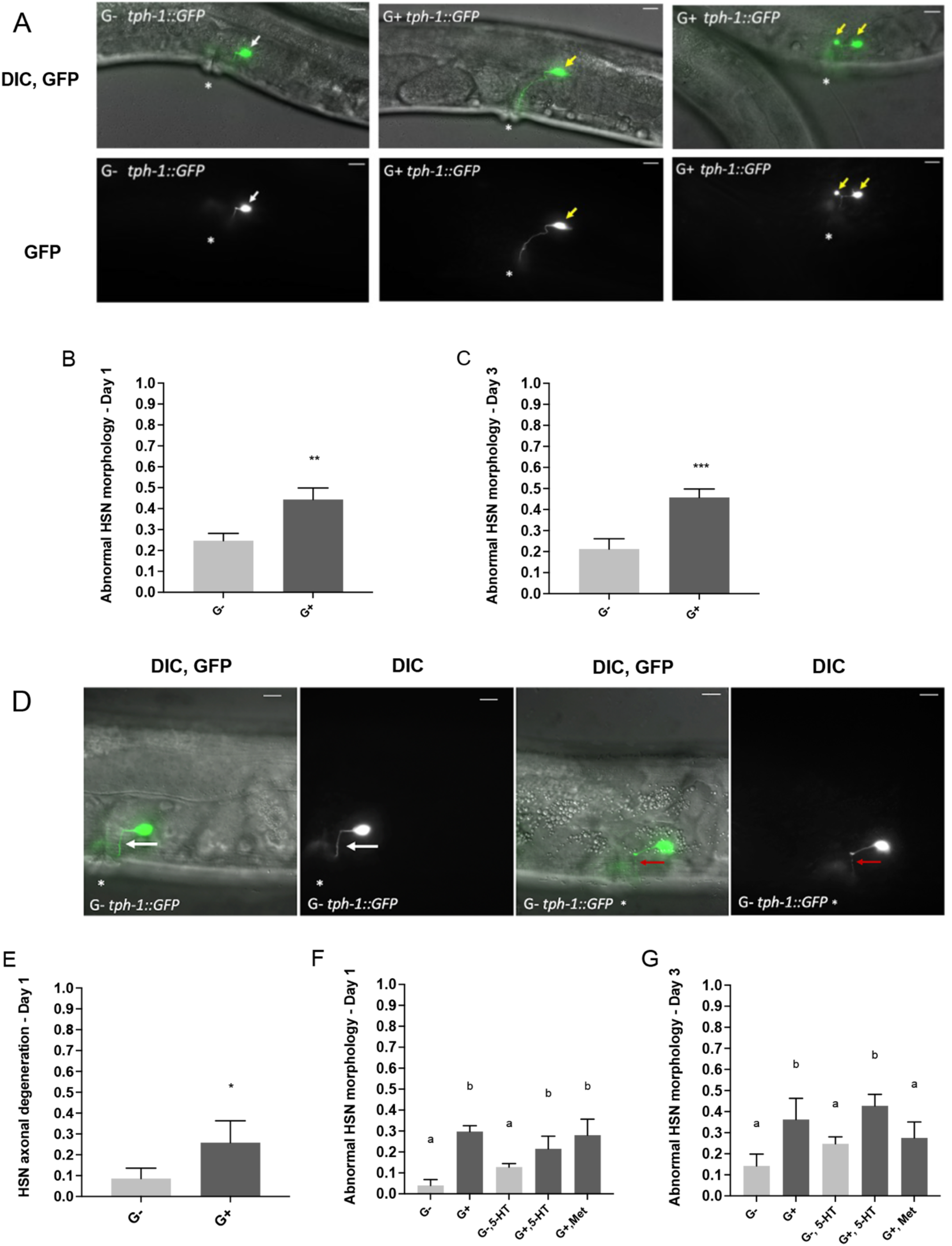
A glucose-supplemented diet induces abnormal morphology and axonal degeneration in the HSN neurons. (A-H) The *tph-1::GFP* reporter was used to assess the HSN neurons in animals fed a standard OP50 diet (G−) or glucose-supplemented diet (G+). (A) Shown are representative animals fed a standard OP50 or glucose-supplemented diet. The upper panel is the merged DIC and fluorescent image and the lower panel is the fluorescent micrograph. For all images, left is anterior, right is posterior, and * indicates location of the vulva. White arrows point at an HSN with normal morphology and yellow arrows point at an HSN with abnormal morphology. Bar = 10 µm. (B-C) The proportion of animals with an abnormal HSN morphology phenotype was assessed in day-1 old or day-3 old *tph-1::GFP* hermaphrodites fed a control (G−) or glucose-supplemented diet (G+). The ** indicates p < 0.01; The *** indicates p < 0.001. (two-tailed unpaired t-test; at least three independent experiments were conducted with n>100). (D) The *tph-1::GFP* reporter was used to assess HSN axonal degeneration in animals fed a standard OP50 diet (G−) or glucose-supplemented diet (G+). The upper panel is the merged DIC and fluorescent image and the lower panel is the fluorescent micrograph. For all images, left is anterior, right is posterior, and * indicates location of the vulva. White arrow points at normal HSN axon. Red arrow points at HSN with axonal degeneration. Bar = 10 µm. (E-F) The proportion of animals with HSN axonal degeneration was assessed in day-1 or day-3 adults *tph-1::GFP* hermaphrodites fed a control (G−) or glucose-supplemented diet (G+). (E) The * indicates p < 0.05; (two-tailed unpaired t-test; at least three independent experiments were conducted with n>100). (F) The ** indicates p < 0.01 (two-tailed unpaired t-test; at least three independent experiments were conducted with n>100). (G−H) The proportion of animals with HSN axonal degeneration was assessed in day-1 or day-3 adult *tph-1::GFP* hermaphrodites fed a glucose diet (G+) and methiothepin (1.5 µM) or standard diet with serotonin 5-HT (56 µM). (G−H) Identical letters represent groups with no significant difference. Non-identical letters represent groups with significant difference (p value < 0.01; one-way ANOVA, Tukey’s multiple comparisons; at least three independent experiments were conducted with n >100).

We examined if glucose had an indirect role in HSN abnormalities. To test if it was the holding of eggs within the uterus that induced HSN abnormality, we used floxuridine (5’-fluorodeoxyuridine, FUdR) to eliminate germline progression and egg production in wild-type animals fed a glucose-supplemented diet. The glucose-supplemented FUdR-fed animals, absent of eggs within the uterus, had a significant increase in the proportion of abnormal HSN morphology relative to those fed a standard OP50-diet (Figure S3B, C). We next tested if abnormal HSN migration occurred in the animals fed a glucose-supplemented diet. Using the *kal-1::GFP* reporter to track HSN migration (Kennedy *et al*. 2013) we did not observe a significant difference in HSN migration in animals fed a control or glucose diet (Figure S3D). Combined, these data support the idea that glucose, and not the holding of eggs within the uterus or abnormal HSN migration, negatively impacts the HSN morphology and that the IUEH phenotype is not due to an HSN migration issue.

### A glucose-supplemented diet impacts other behaviors and NSM morphology

In addition to having a role in egg-laying behavior, serotonin and dopamine are associated with other behaviors in *C. elegans*. For example, serotonin has a crucial role in regulating the feeding behavior of pharyngeal pumping (Sze *et al*. 2000), olfaction-driven repulsive behavior (e.g. response to 1-nonanol) and dopamine acts as a neuromodulator in chemosensation and influencing downstream behaviors such as locomotion and arousal that follow mechanical stimulation associated with touch response (Mcdonald *et al*. 2007; Kimura *et al*. 2010; Baidya *et al*. 2014). We determined that a glucose-supplemented diet impacts behaviors modulated by these neurotransmitters, including those that occur downstream. First, the pumping rate of animals fed a glucose-supplemented diet increases relative to animals fed a control diet (Figure 7A). Additionally, the touch response decreases in N2 animals fed a glucose-supplemented diet relative to animals fed a control diet (Figure 7B). Finally, N2 animals fed a glucose-supplemented diet exhibit a significant delay in response to 1-nonanol (Figure 7C).

**Figure 7.**
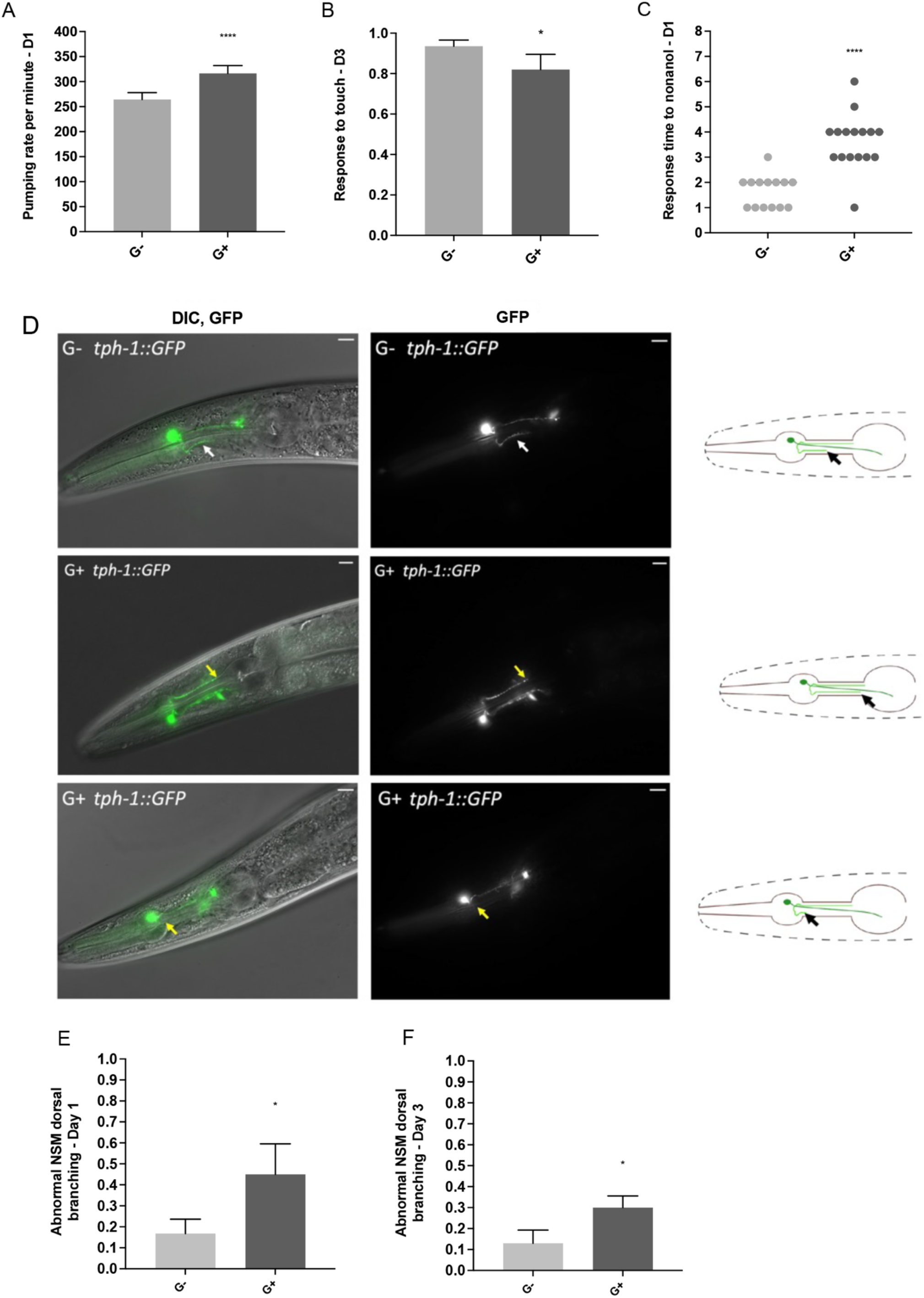
A glucose-supplemented diet impacts serotonin and dopamine associated behaviors and NSM structure. A glucose-supplemented diet impacts (A) pharyngeal pumping in 1-day old N2 hermaphrodites, (B) response to touch in 3-day old N2 hermaphrodites, and (C) response to 1-nonanol in 1-day old N2 hermaphrodites. The **** indicates p <0.0001 and * indicates p < 0.05 (two-tailed unpaired t-test; at least three independent experiments were conducted with n>10). (D) The *tph-1::GFP* reporter was used to assess the NSM dorsal branch in animals fed a standard OP50 diet (G−) or glucose-supplemented diet (G+). Shown are representative animals fed a standard OP50 (G−) or glucose-supplemented diet (G+). The left panels are the merged DIC and fluorescent images and middle panel is the fluorescent micrograph. For all images, left is anterior and right is posterior. Single white arrow points at the end of the NSM dorsal branch. Single yellow arrow points to abnormally long NSM dorsal branch (middle panel) or missing NSM dorsal branch (bottom panel) in animals fed a glucose diet (G+). The right panel is a schematic representation of the NSM dorsal branch. Single black arrow points to the end of the NSM dorsal branch in normal (top panel), abnormally long (middle panel), or missing (bottom panel). Bar = 10 µm (E-F) The proportion of animals with abnormal NSM dorsal branching was assessed in day-1 old or day-3 old *tph-1::GFP* hermaphrodites fed a control (G−) or glucose-supplemented diet (G+). The * indicates p <0.05 (two-tailed unpaired t-test; at least three independent experiments were conducted with n> 100).

The neurosecretory motor (NSM) neuron can be visualized with the *tph-1::GFP* transcriptional reporter and has two branches, posterior and dorsal, that originate from the body of NSM neurons (Axang *et al*. 2008). A normal dorsal branch ends halfway through the isthmus as shown in Figure 7D (white arrows in top panel and black arrows in top panel of the illustrative diagram). The glucose-fed animals exhibit NSM neuron abnormalities (Figure 7D). The NSM abnormality observed includes NSM dorsal branch that is missing or has an abnormal length as it either did not reach the mid region of the isthmus or extended beyond. (Figure 7D yellow arrows in mid and lower panels and black arrows in mid and lower panels in illustrative diagram). Adult hermaphrodites fed a glucose-supplemented diet to 1-day old and 3-day old adulthood had a significant increase in abnormal NSM dorsal branching relative to the animals fed a standard OP50 diet (Figure 7E, F).

## DISCUSSION

Our findings suggest that a high-glucose diet perturbs serotonin signaling, revealing how diet can distort neuromodulatory regulation of egg-laying behavior. Although serotonin typically promotes egg-laying in *C. elegans*, exogenous serotonin administration failed to rescue the glucose-induced IUEH phenotype in wild-type animals and instead exacerbated it. This paradoxical effect suggests that under high-glucose conditions, serotonin signaling may be reprogrammed to drive rather than relieve egg-laying dysfunction. Consistent with this interpretation, IUEH was suppressed in *tph-1* mutants lacking serotonin and by pharmacological inhibition of serotonergic signaling using receptor antagonists such as methiothepin. We also observed transcriptional upregulation of *tph-1* in glucose-fed animals, supporting the interpretation that excessive or misregulated serotonin signaling, rather than its absence, contributes to the glucose-induced IUEH phenotype. Additionally, we demonstrate that a glucose diet affects egg-laying rate, consistent with prior studies (Teshiba *et al*. 2016; Alcantar-Fernandez *et al*. 2018). This insight highlights the potential protective role of dampened serotonergic signaling in response to glucose-induced metabolic stress and suggests a reconfiguration of canonical neuromodulatory functions in altered dietary environments.

Further complexity is revealed through our analysis of serotonin receptor mutants. Glucose-induced IUEH persisted in *ser-1* and *ser-7* mutants, and the phenotype was not suppressed by methiothepin in *ser-1* animals, suggesting that neither receptor alone mediates the glucose–serotonin interaction driving IUEH. Interestingly, *ser-7* mutants displayed IUEH independent of glucose, and combining glucose with exogenous serotonin further exacerbated the phenotype. These results suggest that an uncharacterized serotonin receptor or a downstream serotonin-modulated mechanism is responsible for this effect. Furthermore, the suppression of the glucose-induced IUEH phenotype by chlorpromazine raises the possibility of an intersection between serotonin and dopamine under high-glucose conditions.

Beyond behavioral phenotypes, we also observed structural changes in the serotonergic nervous system. A glucose-supplemented diet disrupted the morphology of the HSN and NSM neurons, including axonal degeneration in the HSN. These findings are particularly notable, given the established link between hyperglycemia and neuropathy in humans; however, the direct impact of chronic hyperglycemia on neuronal structure remains poorly understood. Our results suggest that excess dietary glucose compromises both neuromodulatory signaling and neuronal function and integrity.

In summary, this work establishes a functional and anatomical link between diet and serotonergic modulation of egg-laying in *C. elegans*. This data supports a model in which glucose increases the transcription of *tph-1,* potentially elevating serotonin synthesis and signaling, leading to maladaptive egg-laying dynamics through altered receptor pathways and neuronal integrity. All together, these findings underscore the importance of investigating classical neurotransmitter systems within metabolically perturbed contexts and position *C. elegans* as a powerful model for exploring the impact of high-glucose diet-induced stress on neuronal function and structure.

## MATERIALS AND METHODS

### Strains and growth conditions

*C. elegans* wild-type Bristol strain (N2) and mutant strains were grown on nematode growth media (NGM) plates seeded with *E. coli* OP50 and raised and maintained at 20°C (Brenner 1974). The following genetic strains were used for these studies: *ser-1(ok345)* (Xiao *et al*. 2006), *ser-7(tm1325);ser-1(ok345)* (Hapiak *et al*. 2009), *ser7(tm1325)* (Hobson *et al*. 2006), *bas-1(tm351)* (Hare and Loer 2004), *cat-2(n4547)* (Omura *et al*. 2012), *tph-1p::GFP* (SK4013) (Clark and Chiu 2003), *ida-1p::GFP* (BL5717) (Zahn *et al*. 2001), and *kal-1p::GFP* (Bulow *et al*. 2002).

### Intrauterine egg-hatching, egg-laying rate, and brood size assays

To determine the impact a glucose-supplemented diet had on offspring production intrauterine egg-hatching (which includes matricide events), egg-laying rate, and brood size was determined for hermaphrodites fed either a control diet of NGM with OP50 (noted as G−), or NGM OP50 media supplemented with 27.75 mM (0.5%) glucose (noted as G+). For each respective experiment, the stage of development on the respective diet is noted.

For the intrauterine egg hatching assay, 1-day old adult hermaphrodites were placed on newly seeded NGM OP50 media and allowed to lay eggs for 1-2 hours. The eggs were moved to either an OP50 control diet (G−) or OP50 glucose-supplemented diet (G+) and allowed to reach adulthood. Populations that only contained hermaphrodites (no males) were included in these studies. The adult hermaphrodites were assayed daily to determine if there were one or more hatched embryos, as indicated by larvae within the uterus. These animals, whether alive or dead due to matricide, were scored as having the intrauterine egg hatching phenotype (noted as IUEH throughout this manuscript). The number of animals that exhibited a matricide phenotype was noted and included in the IUEH phenotype for consistency and simplicity. Daily, the surviving animals were transferred to the new respective diet (G− or G+) and the animals were assayed daily for 10 days or if indicated throughout the lifespan.

To determine egg-laying rate, embryos were allowed to hatch and develop on the respective media (G− or G+). The one-day old animals were placed individually onto small NGM plates (5 mL) seeded with the respective media (G− or G+) and allowed to lay eggs for 1 hour and the total number of laid embryos per hour was determined.

To quantify brood-size, embryos were allowed to hatch and develop to the L4 larvae stage on the noted respective media (G− or G+). When the synchronized population reached the L4 larvae stage, as determined by formation of the vulva patch and a bend in the germline, individual hermaphrodites were placed onto a small (5 ml) NGM plates seeded with the respective diet fed during development (G− or G+). The animal was transferred every 24 hours to a new plate containing the same diet and transfer continued until the animal no longer produced offspring. The number of offspring produced daily was quantified to determine brood size.

### Drug Supplementation

The delivery of exogenous serotonin hydrochloride (Fisher Scientific), dopamine (Fisher Scientific), methiothepin mesylate salt (Sigma-Aldrich) or chlorpromazine hydrochloride (Sigma-Aldrich) was adapted from previous studies, respectively (Zarse and Ristow 2008; Baidya *et al*. 2014; Cruz-Corchado *et al*. 2020). The molecules were added to the NGM agar plates (10 mL,100 mm x 15 mm) seeded with OP50 (G−) or OP50 supplemented with glucose (G+) to obtain the noted final concentrations for each respective experiment (56 µM serotonin, 56 µM dopamine, 1.5 µM methiothepin, or 1.5 µM chlorpromazine).

### Fluorescent Microscopy of HSN and NSM

Neuron morphology was assessed within the 1, 3 or 5-day old adult hermaphrodite fed a control or glucose-supplemented diet using a Zeiss Axioscope fluorescent microscope with differential interference contrast (DIC) capabilities (King *et al*. 2021). Adult hermaphrodites were mounted on a 2% agarose slide containing an anesthetic solution (00.1% levimasole and 0.1% tricane). Images were obtained using the Axiosoftware 4.7.1 software and processed using Adobe Photoshop 2020. The left or right hermaphrodite specific motor neuron (HSN) was assessed using the *tph-1::GFP* or *ida-1::GFP* fluorescent reporter strains and the position of the cell body and neurite morphology was determined. An abnormal HSN phenotype was noted when there was a visible defect with cell body position, cell body morphology, or ectopic neurite. Axons that demonstrated a break were scored as having a degeneration phenotype. The left or right neurosecretory motor neuron (NSM) was assessed using the *tph-1::GFP* fluorescent reporter and abnormal branching was determined by the position of the NSM dorsal branch end. Animals with NSM dorsal branch length that did not reach the mid region of the isthmus or extended beyond it were scored as abnormal (Axang *et al*. 2008; Nelson and Colon-Ramos 2013).

### Phenotype Analysis of Glucose-Fed Animals

Similar to previous methods (Raizen D. 2012), pharyngeal pumping was quantified in N2 adult hermaphrodite (1-day and 3-day old) fed a control or glucose-supplemented diet. Using a dissecting microscope, the number of grinder movements within a 1-minute period was determined in three independent experiments in >30 synchronized animals. To assess the capacity to respond to a repulsive agent, N2 adult hermaphrodites (1-day and 3-day old) were washed with M9 buffer twice and a single animal was placed on a sterile unseeded NGM plate, 2 µL of 1-nonanol was placed near the anterior head region of the worm (within 1 cm) (Sammi *et al*. 2022). Using a stopwatch, the time it took for the animal to move away from the 1-nonanol was determined. Three independent experiments of a total of at least fourteen animals were assayed. Animals that did not move from the area and subsequently died were nulled. To assess an anterior touch response, N2 adult hermaphrodites (1-day and 3-day old), on a seeded NGM plate, were brushed with paintbrush hair across the body behind the pharynx region (Chalfie 2014). Animals that backed away when touched were scored as “responding to touch”. The paintbrush hair was sterilized by dipping it into a 70% ethanol solution and allowed to dry prior to individual animal assessment. Three independent experiments for a total of 60 animals were conducted. To determine if N2 or *tph-1(mg280)* adult hermaphrodites retained embryos, the uterus of the adult hermaphrodite, at the specified stage, was examined for the presence of embryos at a 1.5-fold stage or older.

### Quantitative RT-PCR

Non-gravid young adult hermaphrodites (80 per experiment) were collected for mRNA isolation and RT-PCR experiments. Animals were washed twice using M9 and allowed to settle by gravity to remove bacteria from the sample in PCR tubes; samples in 5 µL of M9 were immediately frozen using liquid nitrogen. RNA isolation was performed as previously described (Ly *et al*. 2015). Briefly, 5 µl of 2X worm lysis buffer (10 mM Tris-HCl pH 8.0 (Sigma–Aldrich), 1% Triton X-100 (Sigma-Aldrich), 1% Tween 20 (Bio-Rad Laboratories), 0.5 mM EDTA (Merck), and 0.4 U proteinase K (NEB) were added to the 5 µl worm sample. The mixture was then incubated in a thermocycler at 65 °C for 10 minutes, followed by 85 °C for 1 minute to inactivate the proteinase K. The complementary DNA was generated using a Maxima H Minus cDNA synthesis kit (Thermo Fisher) according to the provided instruction manual. Quantitative RT-PCR was conducted using a StepOnePlus real-time PCR system (Applied Biosystems) and PowerUpSYBR Green Master Mix (Applied Biosystems) as previously described (King *et al*. 2021). REST software was used to calculate the relative expression levels (Pfaffl *et al*. 2002). The mRNA level of *iscu-1* (Y45F10D.4) was used for normalization (Hoogewijs *et al*. 2008). The average of three independent replicates was used and analyzed using unpaired Student’s T-test. The primers used in this study are: *tph-1* fw: CTCCAGAACCAGACACCGTTC; *tph-1* rv: TCAGACGAGAGACCAAATTCAA; *cat-2* fw AAGACATTGGGCTGATGAGTTT; *cat-2* rv: CGCGTGCATCAATTCTCC; *iscu-1*-fw TCCTGCACAAGTTTGCGTTG; *iscu-1*-rv CCGTTATCGTCGACTCGAATCTG.

### Statistics

The statistical test (one-way ANOVA, Tukey’s multiple comparisons, Two-tailed unpaired T-test) and P-values are noted for each experiment within the appropriate figure legend. All datasets are expressed as mean standard deviation, or as a proportion of animals with the noted phenotype. A minimum of three independent experiments were conducted. Statistical analysis was conducted using GraphPad Prism 9 software.

## Supporting information

Table S1_RNA_Seq

## ACKNOWLEDGMENTS

This work was supported by the grant from the National Institute of Health (R15DK109524) to P.A. Padilla. We thank the Caenorhabditis elegans Genetics Stock Center (CGC), which is funded by the NIH office of Research Infrastructure Programs (P40 OD010440).

## Notes

### Competing Interest Statement

The authors have declared no competing interest.

