## Supplementary figures and images for "Glucose promotes neuron morphology abnormalities and egg-laying defects dependent of the serotonin signaling pathway in *Caenorhabditis elegans*"

### Figure S1-3

Figure S1

A

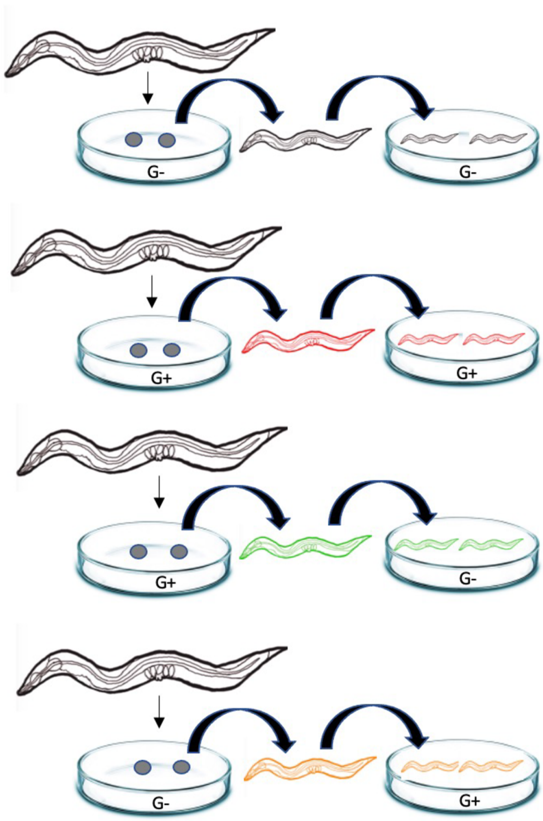

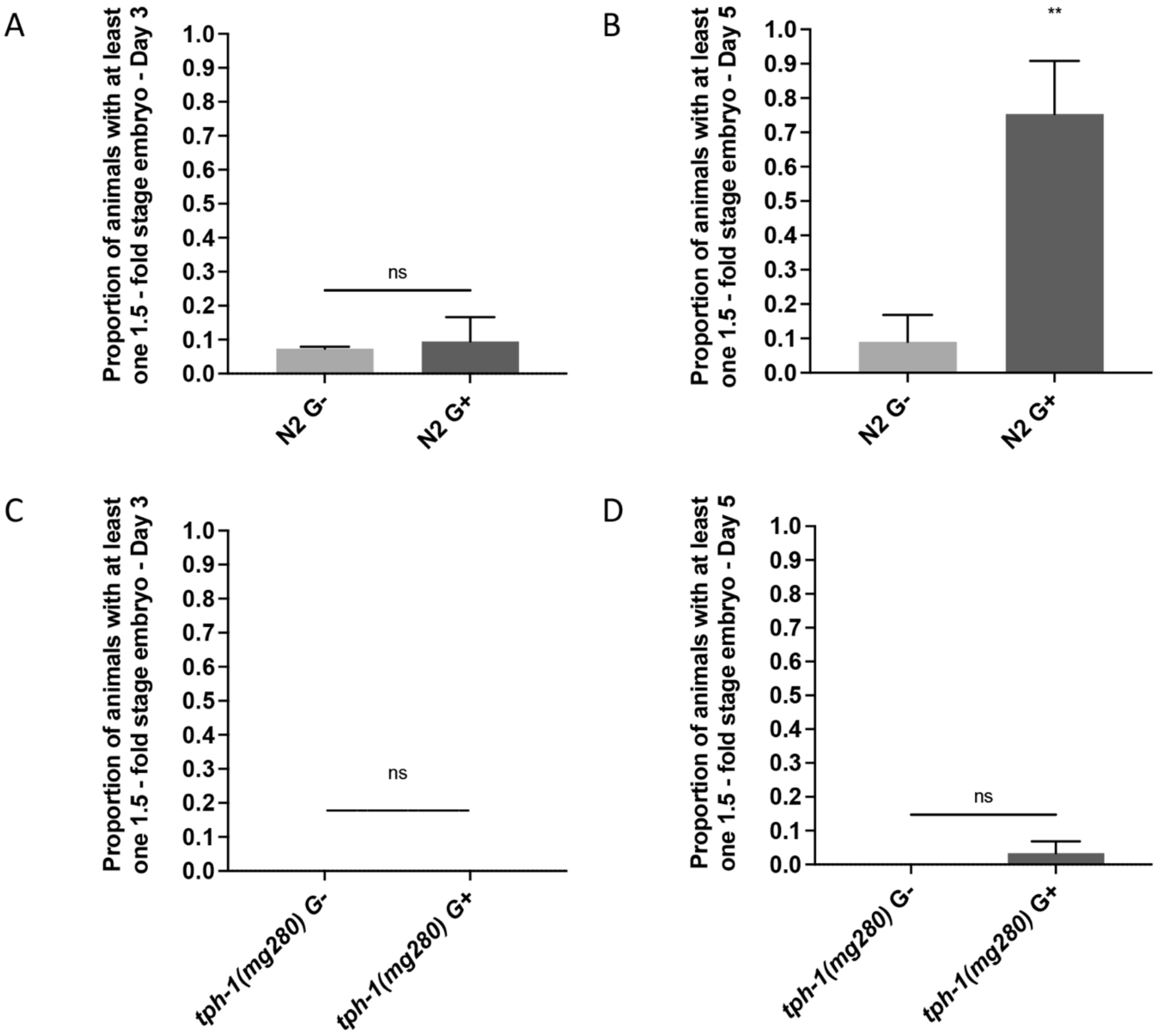

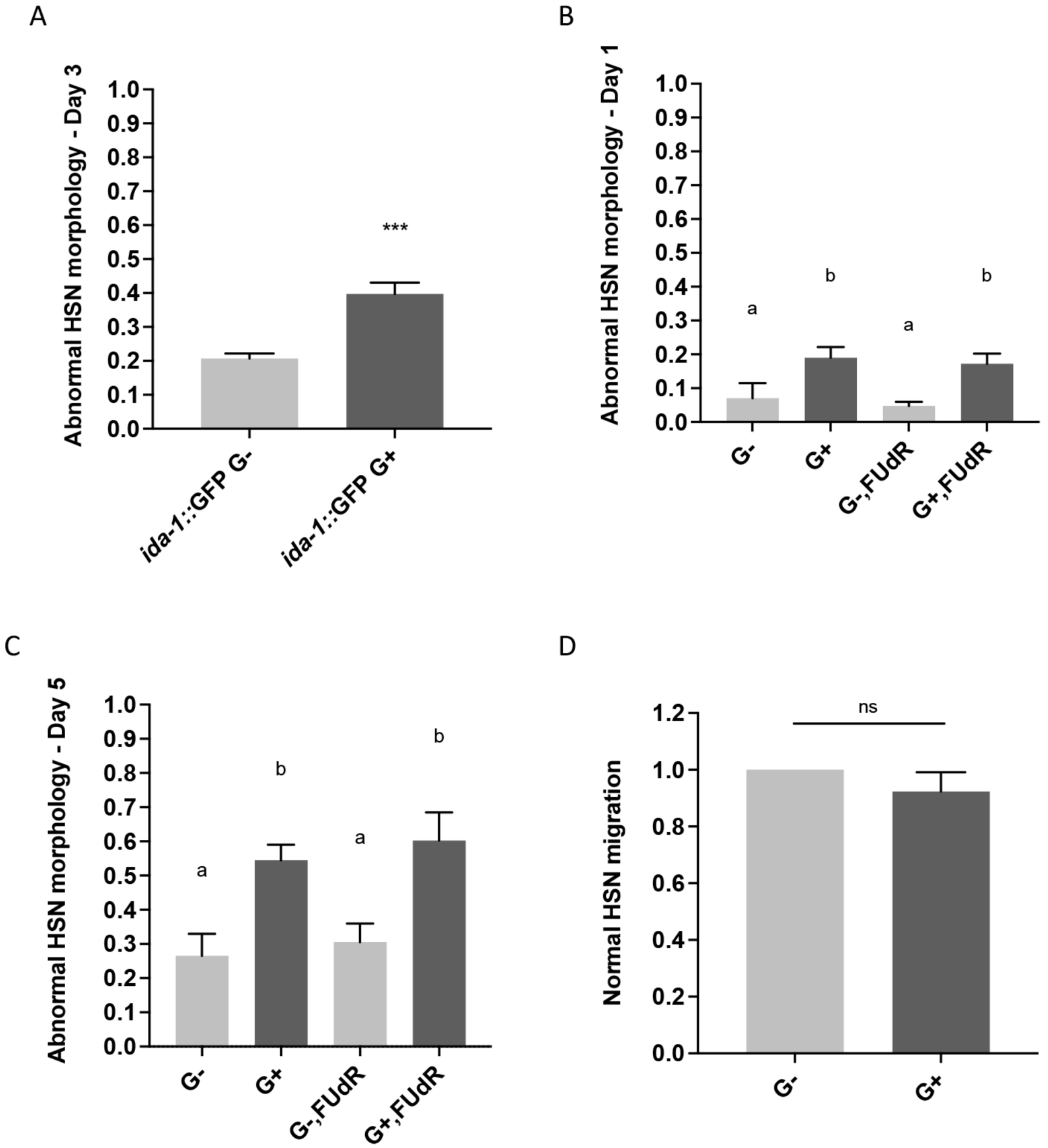
